# *STT3A* is required for recognition of pathogen-derived sphingolipids in Arabidopsis

**DOI:** 10.1101/2025.05.18.654772

**Authors:** Seowon Choi, Motoki Shimizu, Akira Abe, Nobuaki Ishihama, Yuko Ishikawa, Daigo Takemoto, Ken Shirasu, Yoshitaka Takano, Ryohei Terauchi, Hiroaki Kato

**Author notes:** Corresponding author: Hiroaki Kato.

## Abstract

Plants recognize pathogen-associated molecular patterns (PAMPs) via pattern recognition receptors, leading to the activation of pattern-triggered immunity in response to pathogen attack. *<u>P</u>hytophthora infestans* <u>cer</u>amide <u>D</u> (Pi-Cer D) is a sphingolipid from the oomycete pathogen *P. infestans*. Pi-Cer D is cleaved by the plant extracellular ceramidase NEUTRAL CERAMIDASE 2 (NCER2), and the resulting 9-methyl-branched sphingoid base is recognized by the plant receptor RESISTANT TO DFPM-INHIBITION OF ABSCISIC ACID SIGNALING 2 (RDA2) at the plasma membrane to transduce a defense signal. However, additional components are likely involved in sphingolipid recognition, which remain to be identified. Here, we employed a screen based on Lumi-Map technology to look for Arabidopsis (*Arabidopsis thaliana*) mutants with altered defense responses to Pi-Cer D. We identified three mutants showing diminished responses to Pi-Cer D and elf18, each carrying mutations in *STAUROSPORIN AND TEMPERATURE SENSITIVE 3-LIKE A* (*STT3A*), which encodes an oligosaccharyltransferase. The *stt3a* mutants exhibited higher susceptibility to the pathogen *Colletotrichum higginsianum* than the wild type and displayed alterations in NCER2 protein modifications. These findings suggest that STT3A contributes to plant immunity via post-translational modification of proteins including NCER2.

## INTRODUCTION

Plants have evolved complex immune systems that detect and respond to pathogens in the environment (Dodds et al., 2024). Plant immune responses begin with the recognition of pathogen-associated molecular patterns (PAMPs); these conserved molecules are found in a wide variety of microbes, including bacteria, fungi, and viruses (Nejat and Mantri, 2017; Segonzac and Zipfel, 2011). These PAMPs are recognized by pattern recognition receptors (PRRs), which are typically localized at the plasma membrane and activate pattern-triggered immunity (PTI) (Bigeard et al., 2015). PTI is a critical component of the plant immune system that acts as the first layer of defense, preventing pathogen entry and colonization of plant tissues (Jones and Dangl, 2006). PTI initiates with the perception of a ligand by a PRR, followed by the association of the PRR with co-receptors such as BRI1-ASSOCIATED RECEPTOR KINASE 1 (BAK1) and CHITIN ELICITOR RECEPTOR KINASE 1 (CERK1) (Chinchilla et al., 2009; Miya et al., 2007; Ngou et al., 2022). Despite extensive research into the interactions between PAMPs and PRRs, much about PTI signaling pathways remains unknown.

Some PAMPs are sphingolipids; members of this class of lipids consist of a long-chain sphingoid base backbone linked to a fatty acid through an amide bond at the 2-amino group and to a polar head group at the C-1 position via an ester bond (Heung et al., 2006). The 9-methyl-branched sphingoid base is absent from plants but present in fungi and oomycetes and is thus recognized by plants as a non-self molecular pattern to activate defense responses and deter pathogen attack (Koga et al., 1998). Cerebrosides, a group of glycosphingolipids, also contain a 9-methyl-branched sphingoid base as part of their epitope structure and have been identified as elicitor molecules from the rice blast pathogen *Pyricularia oryzae* (Koga et al., 1998; Umemura et al., 2000). Sphingolipids are important for membrane structure and as signaling molecules in both microbes and plants. Although sphingolipids in pathogenic fungi are known to play important roles in the initiation and development of infections afflicting humans, the metabolism and function of sphingolipids from plant pathogenic fungi remain to be fully explored (Zhu et al., 2023).

*Phytophthora infestans* ceramide D (Pi-Cer D) is a PAMP sphingolipid purified from the oomycete *Phytophthora infestans* that promotes resistance in plants (Monjil et al., 2024). In Arabidopsis (*Arabidopsis thaliana*), Pi-Cer D is cleaved by the apoplastic ceramidase NEUTRAL CERAMIDASE 2 (NCER2), yielding a 9-methyl-branched sphingoid base that is recognized by the plasma-membrane-localized lectin receptor-like kinase RESISTANT TO DFPM-INHIBITION OF ABSCISIC ACID SIGNALING 2 (RDA2) (Kato et al., 2022).

However, further investigation is necessary to identify and characterize other factors involved in this recognition, including additional receptors, downstream signaling components, and the regulatory networks that modulate sphingolipid-triggered immune responses.

Here, we used Lumi-Map (Kato et al., 2020) to identify genes involved in sphingolipid recognition. We identified *STAUROSPORIN AND TEMPERATURE SENSITIVE 3-LIKE A* (*STT3A*) as an important component of Pi-Cer D recognition-mediated defense signaling and established its function in the sphingolipid response and pathogen resistance in Arabidopsis.

## RESULTS

### Mutants showing low responses to Pi-Cer D were identified

The expression of the Arabidopsis defense gene *WRKY33* is induced by diverse PAMPs, including the peptide flagellin 22 (flg22) and the sphingolipid Pi-Cer D (Denoux et al. 2008; Kato et al. 2022; Zheng et al. 2006). Therefore, we used an Arabidopsis reporter line harboring the firefly luciferase gene (*LUC*) under the control of the *WRKY33* promoter (p*WRKY33-LUC*) (Kato et al., 2022). We mutagenized the p*WRKY33-LUC* reporter line (the wild type, WT) with ethyl methane sulfonate and screened 10,000 M_2_ seedlings in a previous study (Kato et al., 2022). To select mutants for further analysis, we first confirmed the bioluminescence phenotype in response to Pi-Cer D treatment using M_3_ lines and subsequently classified them based on their responses to other elicitors. Among the mutants, L-08, L-13, and L-51 showed low bioluminescence responses to Pi-Cer D treatment: L-08 and L-51 exhibited minimal responses to Pi-Cer D, whereas L-13 showed a response that was approximately half that observed in WT at its peak (Fig. 1A). We performed reverse-transcription quantitative PCR (RT-qPCR) to evaluate *WRKY33* expression levels in WT and mutant seedlings at 0 and 1 h of treatment with Pi-Cer D. The relative transcript accumulation of *WRKY33* was lower in all three mutants compared to WT specifically following Pi-Cer D treatment (Fig. 1B).

**Figure 1.**
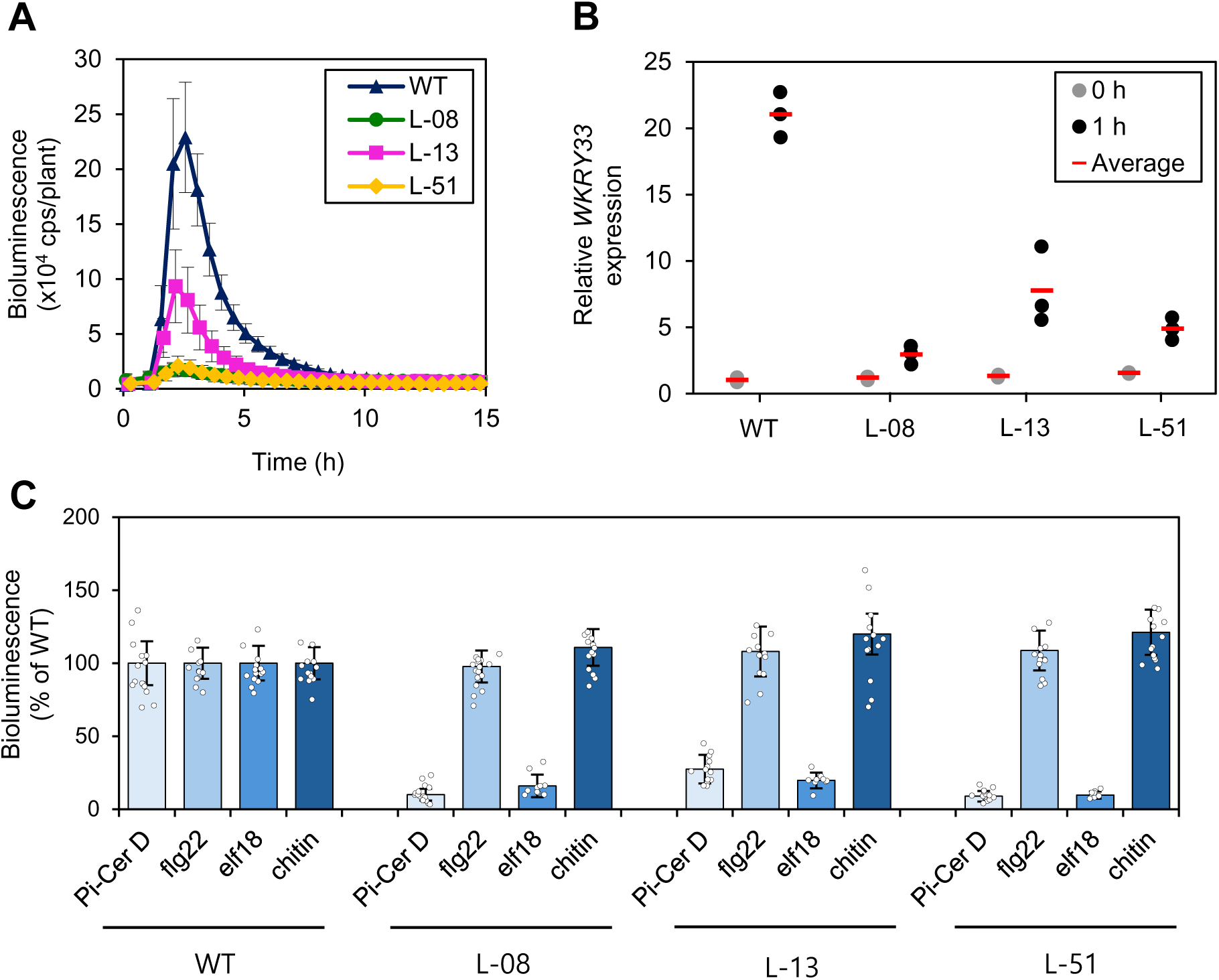
Classification of mutants with altered responses to Pi-Cer D treatment. (A) Bioluminescence patterns of the three identified mutants under Pi-Cer D treatment (means ± SE). Seven-day-old wild type (p*WRKY33-LUC* reporter line; WT) and L-08, L-13, and L-51 mutant seedlings were treated with 0.17 µM Pi-Cer D. The bioluminescence of each seedling was monitored at the indicated time points starting at the onset of treatment using a continuous bioluminescence monitoring system. (B) Relative *WRKY33* expression levels in response to Pi-Cer D treatment in WT and the three mutants, as determined by RT-qPCR. *WRKY33* expression levels were measured in 7-day-old seedlings treated with 0.17 µM Pi-Cer D. (C) Bioluminescence responses of WT and the three mutants following treatment with one of four elicitors (means ± SE). Seven-day-old seedlings were treated with 0.17 µM Pi-Cer D, 0.1 µM flg22, 0.1 µM elf18, or 20 µg/ml chitin, and the bioluminescence of each seedling was monitored. The bioluminescence level in WT was set to 100%, to which the bioluminescence levels in the mutants were normalized.

To classify the mutants showing low bioluminescence responses to Pi-Cer D treatment, we investigated the responses of these mutants to three additional PAMPs: flg22, elf18, and chitin (Fig. 1C). Flg22, a peptide derived from bacterial flagellin, elicits defense responses and is recognized by the receptor FLS2 (Chinchilla et al., 2006). Elf18 is the N-terminal fragment of the bacterial elongation factor Tu recognized by the EF-TU RECEPTOR (EFR) (Zipfel et al., 2006). Chitin is a component of the fungal cell wall that is recognized by the receptor CERK1 (Miya et al. 2007). All three mutants, L-08, L-13, and L-51, exhibited diminished induction of bioluminescence in response to treatment with Pi-Cer D or elf18 compared to WT, whereas their responses to flg22 and chitin were similar to those of WT (Fig. 1C). Therefore, we focused on these three mutants for further analysis.

We also examined the responses of the three mutants to the 9-methyl-branched sphingoid base (4E,8E)-9-methyl-4,8-sphingadienine (9Me-Spd), nlp24, 3-hydroxydecanoic acid (3-OH-FA), and cello-oligosaccharides (COS) (Supplementary Fig. S1). Nlp24 is a peptide derived from necrosis-and ethylene-inducing peptide 1-like proteins (Albert et al., 2015). 3-OH-FA is perceived by the lectin receptor kinase LIPOOLIGOSACCHARIDE-SPECIFIC REDUCED ELICITATION (LORE) (Kutschera et al., 2019). COS are small carbohydrate molecules derived from the partial hydrolysis of cellulose that can elicit a broad-spectrum immune response against several pathogens (Chen et al., 2021; Kongala and Kondreddy, 2023). The mutants showed lower induction of bioluminescence in response to treatment with 9Me-Spd or COS compared to WT, whereas they had normal responses to treatment with nlp24 or 3-OH-FA (Supplementary Fig. S1). The similar responses to various elicitors exhibited by the three mutants suggest that the causative genes of the three mutants are likely involved in the same pathway related to plant responses against Pi-Cer D, 9Me-Spd, elf18, and COS.

### Mutation of *STT3A* results in the low response to Pi-Cer D

To identify the causal mutations of L-08, L-13, and L-51, we crossed each mutant line to the p*WRKY33-LUC* reporter line (the parental line of the mutants) to produce F_1_ progeny, which were self-pollinated to obtain F_2_ seeds. We then tested the F_2_ segregating population derived from each mutant using Lumi-Map (Kato et al. 2020) to select F_2_ seedlings with the mutant phenotype of a reduced bioluminescence response to Pi-Cer D treatment. Following Pi-Cer D treatment, we chose 30 F_2_ individuals showing low bioluminescence responses for genomic DNA extraction. For each F_2_ population, we obtained equal amounts of leaf material from each individual showing a mutant phenotype, mixed the materials together, and extracted DNA from this mixture. We sequenced the DNA samples from each mutant line to detect single nucleotide polymorphisms (SNPs) relative to the p*WRKY33-LUC* reporter line, whose genome was also sequenced. When we plotted the SNP-index as a function of SNP position, we identified a single genomic region with SNP-index values close to 1 for all three mutants. Importantly, the three mutants displayed one large SNP-index peak at the identical position (6 Mb) on chromosome 5 (Fig. 2).

**Figure 2.**
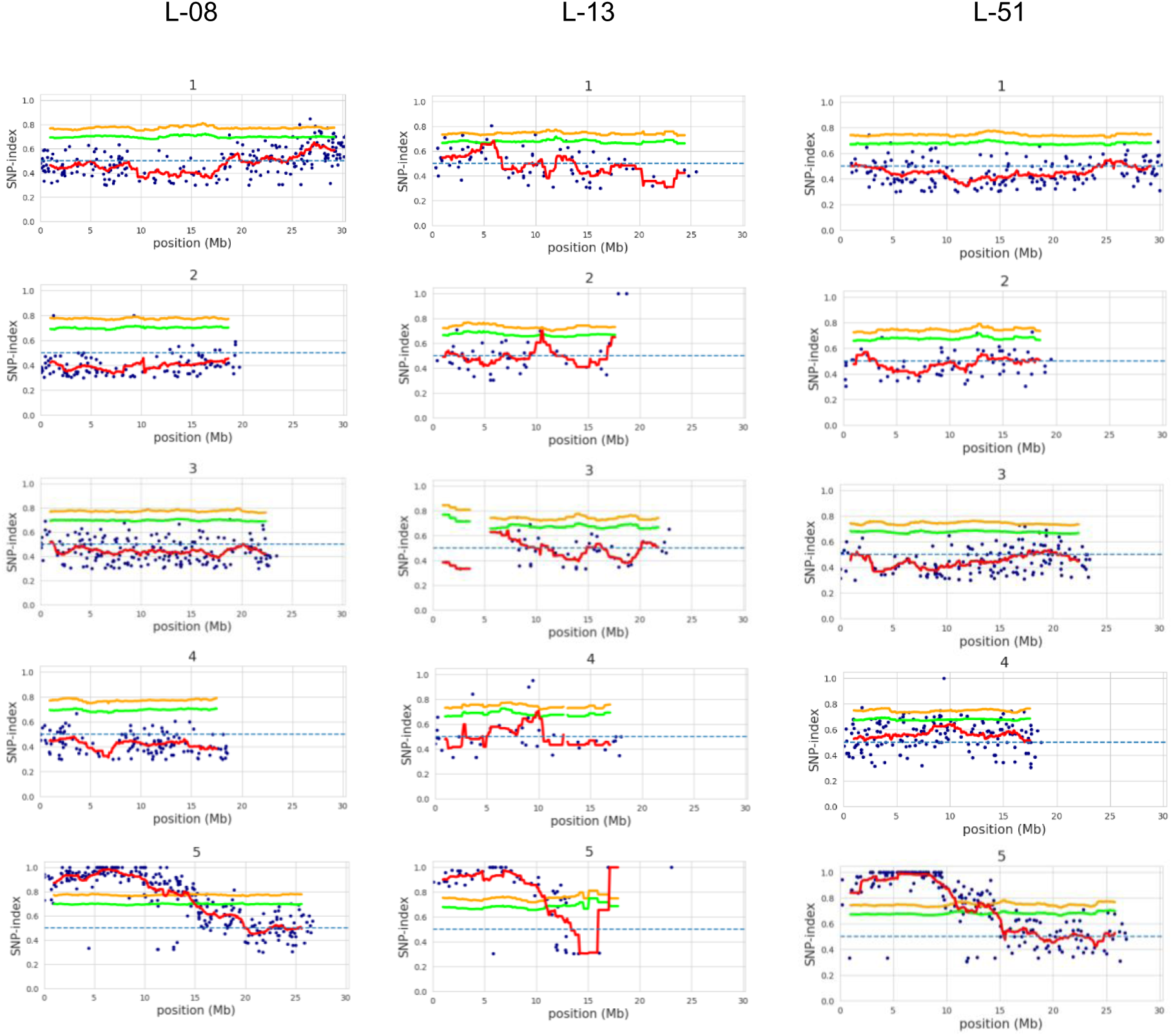
MutMap analysis of the L-08, L-13, and L-51 mutants. The highest SNP-index peak within a genomic region indicates the location of the causative mutation. An SNP-index peak around the 6-Mb region on chromosome 5 was observed in all three mutants: L-08, L-13, and L-51. Blue dots represent individual mutations in each mutant. The red line represents average SNP-index values across a 2-Mb sliding window with 10-kb increments. The green and yellow lines indicate the 95% and 99% confidence limits, respectively, of SNP-index values under the null hypothesis of an SNP-index = 0.5 assuming no linkage between the SNP and the causal mutation.

Based on the p*WRKY33-LUC* reporter line reference genome, the region with the highest SNP-index value in the L-08, L-13, and L-51 mutants contains multiple candidate genes, with the gene *STT3A* harboring mutations in all three mutants (Supplementary Table S1). *STT3A* encodes an oligosaccharyltransferase that plays a pivotal role in the glycosylation of diverse proteins (Cheng et al., 2022). The three mutants harbored different types of mutations: splicing junctions in L-08 and L-13, and an amino acid substitution in L-51 (Fig. 3A and Supplementary Table S1). Genetic complementation of the three mutants with a wild-type genomic *STT3A* fragment led to the recovery of a normal bioluminescence response following Pi-Cer D treatment (Fig. 3B). These results demonstrate that the poor induction of the p*WRKY33-LUC* reporter to Pi-Cer D treatment in the L-08 (*stt3a-3*), L-13 (*stt3a-4*), and L-51 (*stt3a-5*) mutants is caused by mutations in *STT3A*. Furthermore, we obtained the Arabidopsis line *stt3a-2*, with a T-DNA insertion in *STT3A* (Fig. 3A, Supplementary Fig. S2). This mutant showed compromised induction of *WRKY33* expression after treatment with Pi-Cer D, as determined by RT-qPCR (Fig. 3C). These results suggest that *STT3A* is involved in defense signaling mediated by Pi-Cer D in Arabidopsis.

**Figure 3.**
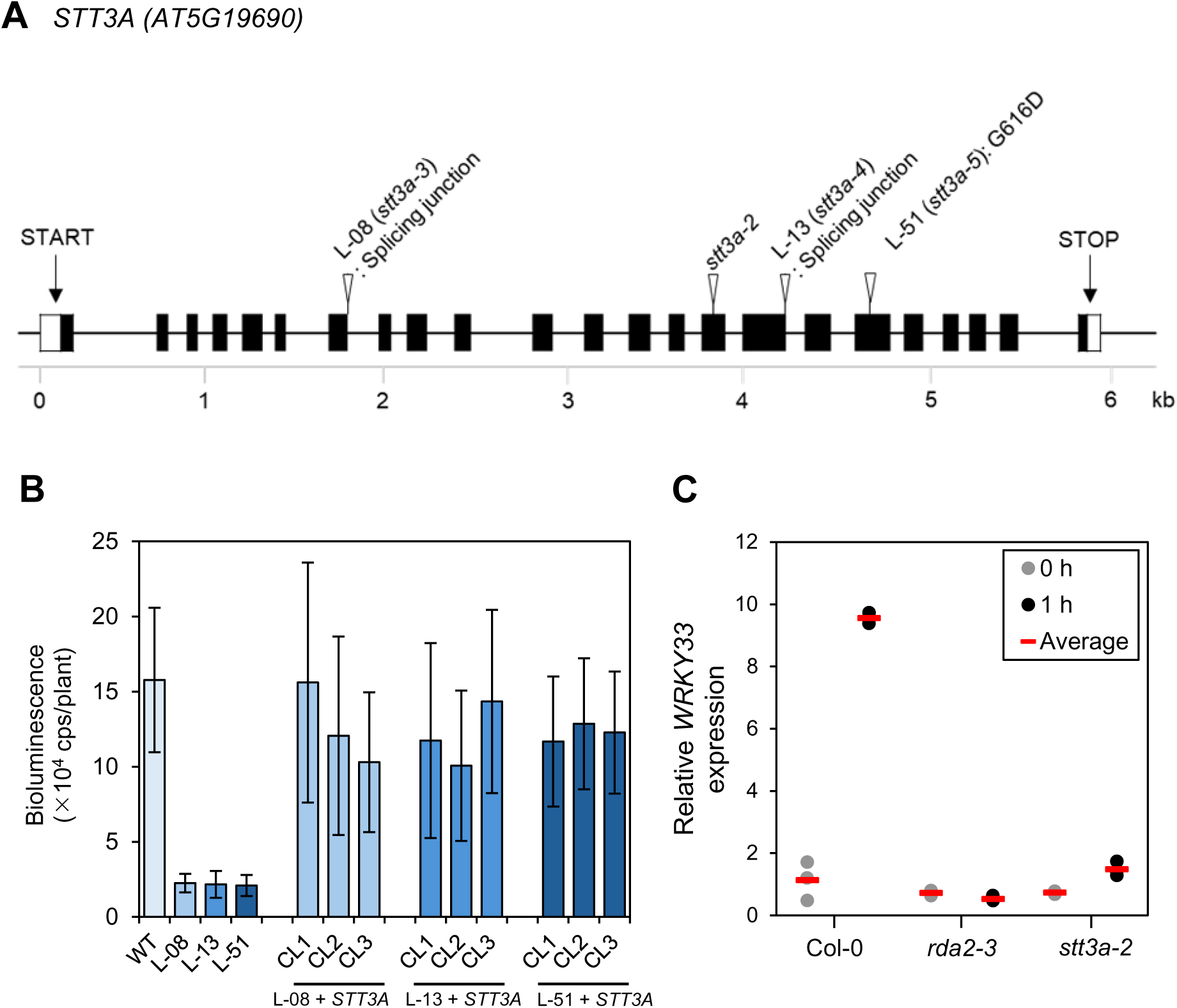
*STT3A* is involved in the recognition of Pi-Cer D in plants. (A) *STT3A* gene structure and the locations of the causative mutations in the three mutants, *stt3a-3* (L-08), *stt3a-4* (L-13), and *stt3a-5* (L-51). Black rectangles indicate exons; lines represent introns. The locations of causative mutations are represented by white triangles, with their mutation types indicated. (B) Genetic complementation of the three mutants with a wild-type copy of the *STT3A* gene. The wild-type *STT3A* gene was introduced into each mutant background. The p*WRKY33-LUC* reporter line (WT), the L-08, L-13, L-51 mutants, and complementation lines for each mutant were treated with 0.17 µM Pi-Cer D (means ± SE); bioluminescence was monitored using a continuous bioluminescence monitoring system. (C) Relative *WRKY33* expression levels in Col-0 and the T-DNA insertion lines *stt3a-2* and *rda2-3*, as determined by RT-qPCR. Seven-day-old seedlings were used for the analysis. The seedlings were sampled at 0, 1, and 3 h of treatment with 0.17 µM Pi-Cer D.

### *STT3A* is required for defense signaling in Arabidopsis following sphingolipid recognition

To determine the role of *STT3A* in defense signaling, we evaluated the expression of defense-related genes and the activation of Mitogen-Activated Protein Kinase (MAPK) signaling following Pi-Cer D treatment of WT, *stt3a-3* (L-08), *stt3a-4* (L-13), and *stt3a-5* (L-51) seedlings. *PENETRATION 2* (*PEN2*) encodes a glycosyl hydrolase that localizes to peroxisomes and acts as a component of an inducible preinvasion resistance mechanism (Lipka et al., 2005). *CYTOCHROME P450 FAMILY 81 SUBFAMILY F2* (*CYP81F2*) is involved in glucosinolate metabolism, and its loss-of-function mutants show impaired resistance to fungal attack (Bednarek et al., 2009; Hunziker et al., 2020). *ETHYLENE RESPONSIVE FACTOR 6* (*ERF6*) encodes a central regulator of stress-induced inhibition of plant growth that is involved in the response to reactive oxygen species (Li et al., 2025; Sewelam et al., 2013). RT-qPCR analysis detected lower expression of these three marker genes in the three *stt3a* mutant lines compared to WT following treatment with Pi-Cer D for 1 h or 3 h (Fig. 4A). The *stt3a-2* T-DNA insertion line also showed compromised induction of *PEN2*, *CYP81F2*, and *ERF6* expression in response to the same treatment (Supplementary Fig. S3A).

**Figure 4.**
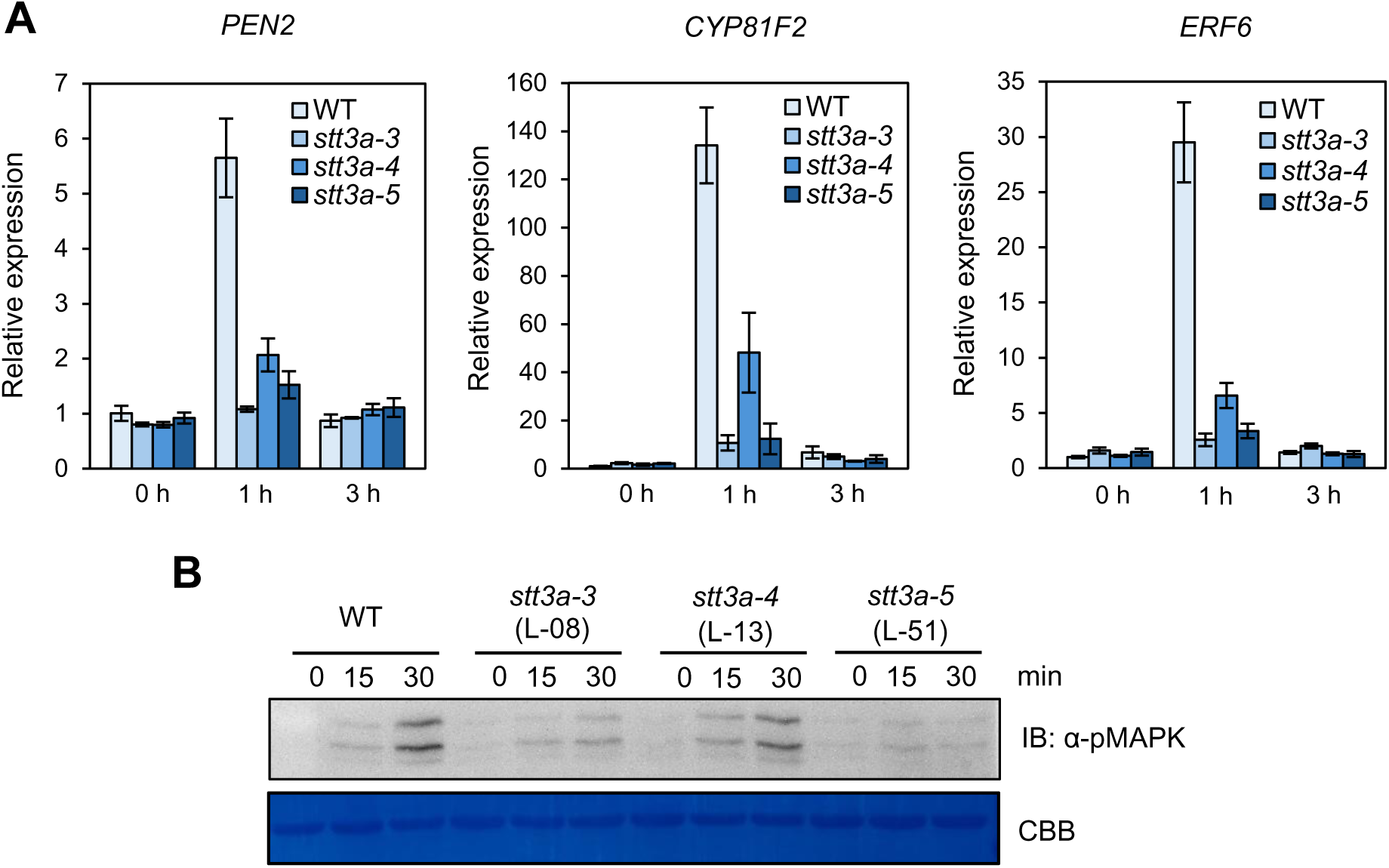
*STT3A* is required for the defense response following the recognition of Pi-Cer D. (A) Relative expression levels of the defense-related genes *PEN2*, *CYP81F2*, and *ERF6* as determined by RT-qPCR (means ± SE). Seven-day-old seedlings of the p*WRKY33-LUC* reporter line (WT), *stt3a-3* (L-08), *stt3a-4* (L-13), and *stt3a-5* (L-51) were treated with 0.17 µM Pi-Cer D. The seedlings were sampled at 0, 1, and 3 h after treatment (*n* = 3). (B) MAPK phosphorylation assay in WT and the three *stt3a* mutants. Seven-day-old seedlings were sampled at 0, 15, or 30 min of treatment with 0.17 µM Pi-Cer D. Phosphorylated MAPKs were visualized with anti-phospho-p44/p42 MAPK antibody. Equal protein loading was determined by staining the membrane with Coomassie brilliant blue (CBB).

We also investigated MAPK activation in WT and the three mutants following the treatment of seedlings with Pi-Cer D (Fig. 4B). WT seedlings exhibited clear MAPK activation at 30 min after the onset of treatment. By contrast, all mutants exhibited diminished MAPK activation. Similarly, the *stt3a-2* T-DNA insertion line showed lower MAPK activation compared to Col-0 (Supplementary Fig. S3B). As a negative control, the *rda2-3* mutant exhibited no induction of marker gene expression and no MAPK activation.

We also observed lower expression of the marker genes and lower MAPK activation when we used 9Me-Spd as the elicitor instead of Pi-Cer D (Supplementary Fig. S4, Supplementary Fig. S5).

### *STT3A* contributes to resistance against the fungal pathogen *Colletotrichum higginsianum*

The lower expression of defense-related genes and the diminished activation of MAPK in the *stt3a* mutants suggested that *STT3A* is required for plant defense against pathogens. To test this hypothesis, we inoculated WT, *stt3a-3* (L-08) *stt3a-5* (L-51), and *pad3-1* (susceptible control; Glazebrook and Ausubel, 1994), which lacks PHYTOALEXIN DEFICIENT 3 function, with the pathogenic fungus *Colletotrichum higginsianum*. The *stt3a-3* and *stt3a-5* mutants were more susceptible to *C. higginsianum* than WT, as evidenced by the size of lesions developing on leaves (Fig. 5). Furthermore, the *stt3a* mutants were more susceptible to *C. higginsianum* than *rda2-5* (Fig. 5). These results suggest that *STT3A* affects the defense of Arabidopsis plants against fungal infection.

**Figure 5.**
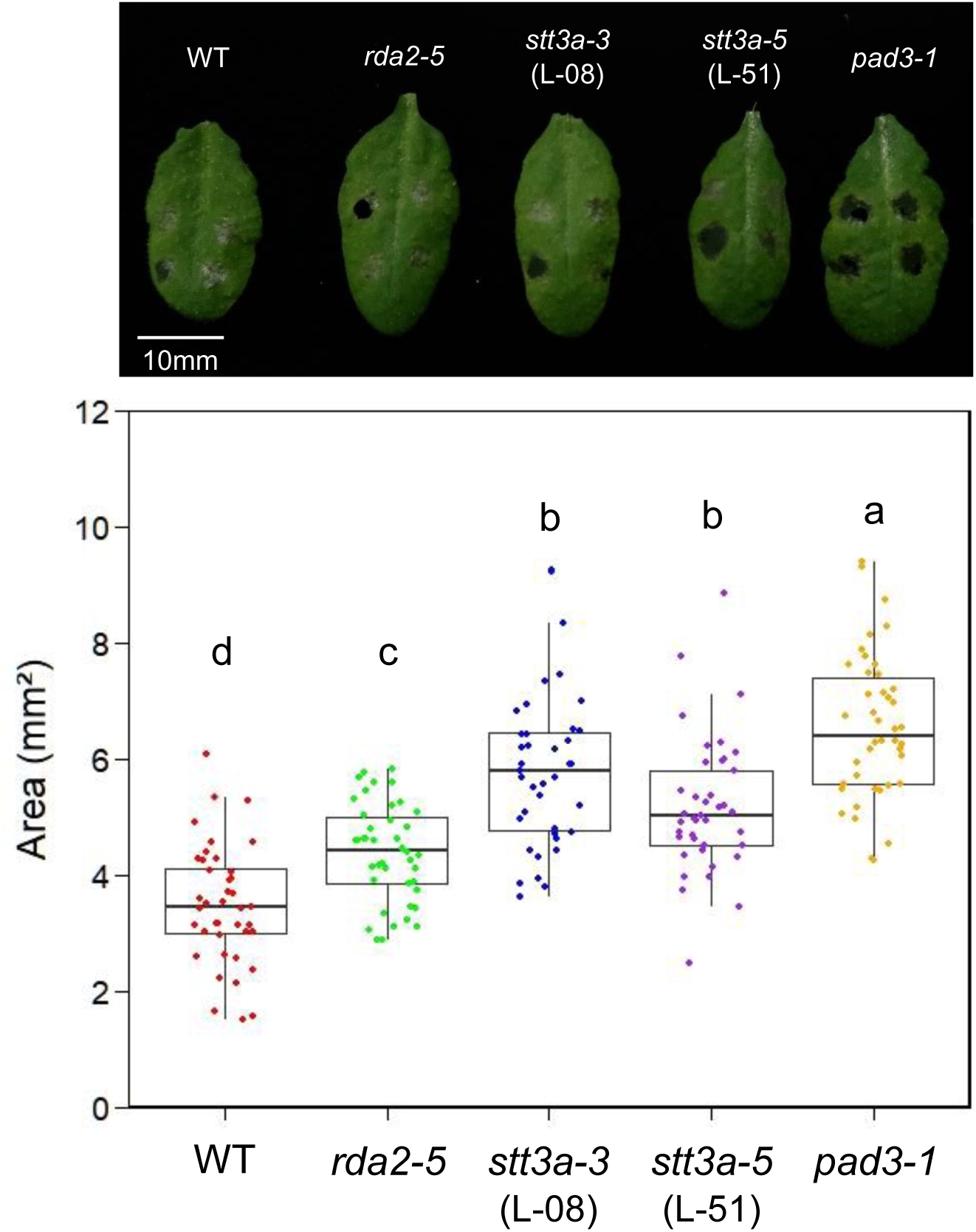
The *stt3a* mutants show enhanced susceptibility to *C. higginsianum*. Six-week-old plants of the p*WRKY33-LUC* reporter line (WT), *rda2-5*, *stt3a-3*, *stt3a-5*, and *pad3-1* were inoculated with a *C. higginsianum* spore suspension containing 1.5 × 10^5^ conidiospores/ml. Top, photograph showing typical lesions on the leaves of WT and the mutants following inoculation with *C. higginsianum*. Bottom, boxplots showing lesion area in WT and the *stt3a* mutants *rda2-5*, and *pad3-1*. Lesion size was measured 4 days after inoculation. Box bounds represent the lesion area, center line represents the median, and whiskers indicate the range of the maximum or minimum data. Different lowercase letters represent statistically significant differences (*p* < 0.05), as revealed by Tukey’s test. The results were replicated in three separate experiments.

### *STT3A* affects the molecular size of NCER2

*STT3A* is involved in the post-translational modification of proteins (Koiwa et al. 2003). We determined that the reduction in the bioluminescence response in the *stt3a* mutants was more pronounced in response to Pi-Cer D vs. 9Me-Spd treatment (Fig. 1C and Supplementary Fig. S1), suggesting that *STT3A* exerts a greater influence on NCER2 than RDA2. The neutral ceramidase of fruit fly (*Drosophila melanogaster*) undergoes glycosylation, a common modification involved in protein secretion and stability (Yoshimura et al., 2022). Therefore, we focused on the abundance and molecular size of NCER2 in the WT and *stt3a* mutants.

NCER2 comprises two distinct structural units; when the N-terminal neutral/alkaline non-lysosomal ceramidase domain is cleaved from the C-terminal portion of the protein, the two resulting fragments form a complex (Kato et al., 2022). We performed immunoblotting with an antibody that recognizes the C-terminus of NCER2 (anti-NCER2) in WT, the *stt3a* mutants (*stt3a-3* and *stt3a-5*), and a *ncer2* mutant (*ncer2-2*; Kato et al., 2022), using total protein extracts and apoplast wash fluid, which should be enriched in secreted proteins (Fig. 6). In WT, we detected a single band (of ∼50 kDa) corresponding to the expected molecular weight of the C-terminus of NCER2, whereas in *ncer2-2*, no band was observed. By contrast, we observed a change in the mobility of NCER2 in *stt3a-3* and *stt3a-5*, indicating a difference in the molecular mass of NCER2. These results suggest that NCER2 is regulated by STT3A-mediated protein modification.

**Figure 6.**
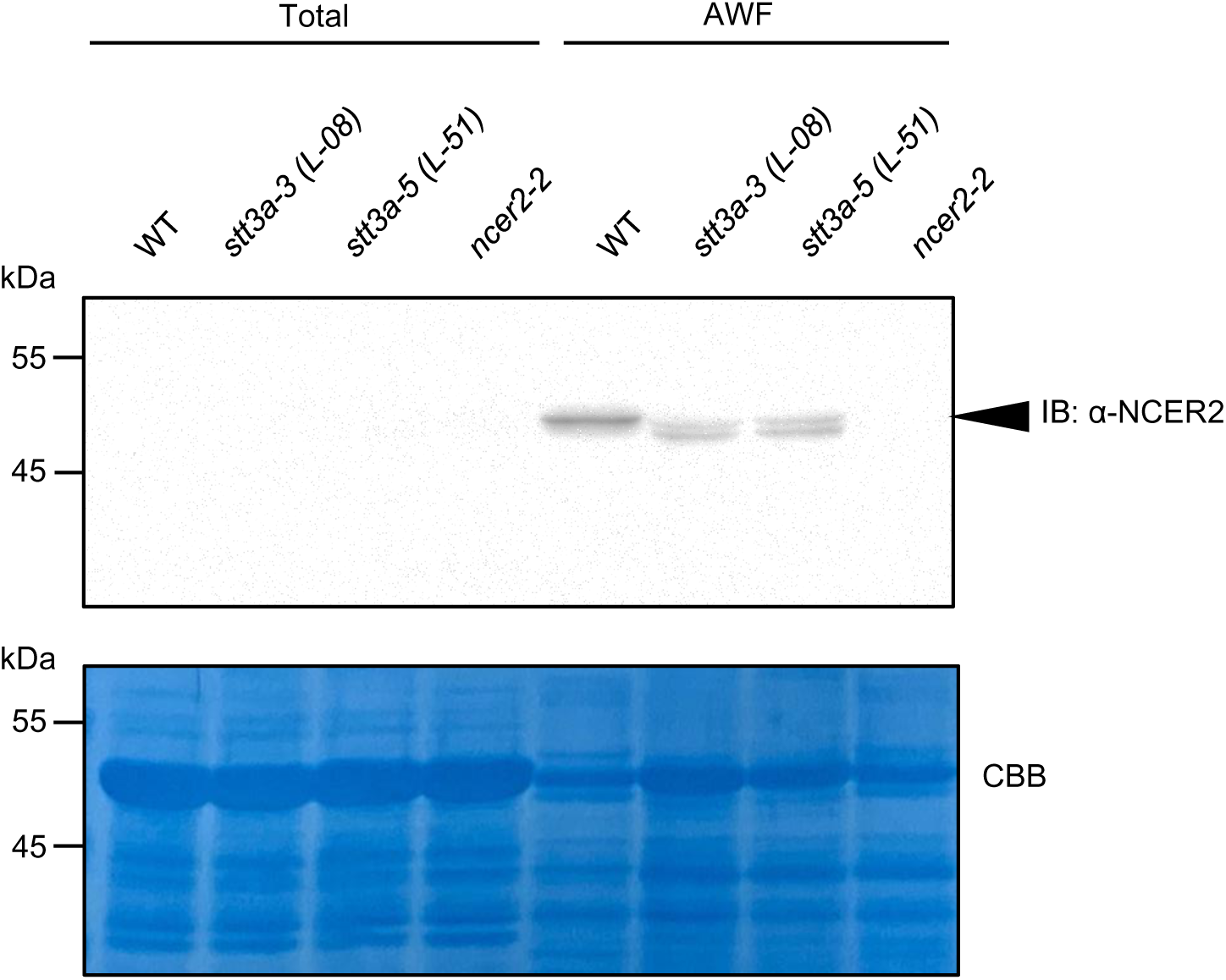
Modification of NCER2 is influenced by *STT3A*. Five-week-old plants were used for immunoblotting (IB) of total proteins and apoplast wash fluid (AWF) in the p*WRKY33-LUC* reporter line (WT) and the *stt3a*-3, *stt3a-5*, and *necr2-2* mutants. NCER2 was detected with a specific antibody against NCER2. Equal protein loading was determined by staining the membrane with CBB. The results were replicated in three separate experiments.

## DISCUSSION

Our Lumi-Map analysis revealed that *STT3A* is required for defense responses in Arabidopsis following the recognition of sphingolipids (Figs. 1–3). *STT3A* is involved in protein N-glycosylation, which supports proper protein folding and function. For example, STT3A-mediated N-glycosylation is important for the function of cellulose biosynthesis enzymes (Kang et al., 2008). In Arabidopsis *stt3a* mutants, defects in N-glycosylation result in insufficient glycosylation of β-glucosidase, an enzyme responsible for converting conjugated forms of the phytohormones abscisic acid and auxin into their active forms (Jiao et al., 2020a). *STT3A* also participates in temperature sensitivity, salinity tolerance, and responses to abiotic stress through increased stomatal density (Jiao et al., 2020b; Liu et al., 2018; Rips et al., 2014; Zhang et al., 2009).

Although all three mutants analyzed harbor mutations in *STT3A*, their phenotypes in reporter assays and their defense responses differed (Fig. 1, Fig. 4, and Supplementary Fig. S4).

The *stt3a-3* and *stt3a-5* mutants showed the weakest defense responses, whereas *stt3a-4* exhibited intermediate defense responses compared to WT (Figs. 1 and 4). Since *stt3a-3* and *stt3a-4* are predicted to affect splicing, *STT3A* transcripts in these two mutants might be different from those of WT. STT3A in *stt3a-5* harbors a G616D mutation that is likely located in the catalytic domain or the dolichol-binding lipid-exposed groove, both of which are essential for oligosaccharyltransferase activity, thus potentially impairing N-glycosylation efficiency (Niu et al., 2020; Wild et al., 2018). Therefore, the mutation in *stt3a-5* might inhibit the enzymatic activity of STT3A or affect its stability.

NCER2 is a ceramidase located in the plant apoplast that converts microbial-derived ceramides into sphingoid bases (Kato et al., 2022). We observed a band shift for C-terminus of NCER2 in apoplast fluid collected from the *stt3a* mutants (Fig. 6), suggesting that NCER2 is post-translationally modified by *STT3A*. Our results suggest that NCER2 is a target of STT3A-mediated N-glycosylation.

From the perspective of plant immunity, STT3A is known to maintain the stability and function of plasma membrane–localized PRRs including EFR and leucine-rich repeat receptor-like kinases (LRR-RLKs) via their N-glycosylation (Häweker et al., 2010; Saijo et al., 2009). In this study, *WRKY33* transcription in the three *stt3a* mutants was also insensitive to treatment with elf18 (Fig. 1), confirming a role for STT3A in EFR regulation (Häweker et al., 2010; Saijo et al., 2009). Moreover, in experiments with various elicitors, the *stt3a* mutants showed less pronounced responses to 9Me-Spd and COS (Supplementary Fig. S1). RDA2 recognizes 9Me-Spd and triggers PTI (Kato et al., 2022). Therefore, RDA2 may also require modification by *STT3A* for its full function. Since we were not able to detect RDA2 in immunoblots, we did not examine its potential modifications but hope to explore this question in the future. Cellulose-derived COS, along with xylooligosaccharides (XOS), are recognized by leucine-rich repeat-malectin receptor kinases such as IMPAIRED GLYCAN PERCEPTION 1 (IGP1, also reported as CELLOOLIGOMER-RECEPTOR KINASE 1 [CORK1]), IGP3, and IGP4 (Fernández-Calvo et al., 2024; Pring et al., 2023; Tseng et al., 2022); these receptors may also be targets of modification by STT3A.

The *stt3a* mutants were less resistant to the fungal pathogen *C. higginsianum* than WT (Fig. 5). *C. higginsianum* contains sphingolipids including 9Me-Spd; their recognition by RDA2 might be important for plant defense against this pathogen. The lower expression of *PEN2* and *CYP81F2* in the *stt3a* mutants following sphingolipid treatment (Fig. 4 and Supplementary Fig. S4) may result in the compromised production and accumulation of antimicrobial compounds, leading to the lower resistance against filamentous fungi (Fig. 5). Notably, the *stt3a* mutants were more susceptible to *C. higginsianum* than the *rda2-5* mutant (Fig. 5). This difference suggests that in addition to signaling mediated by NCER2 and RDA2, multiple pathways for the recognition of PAMPs and damage-associated molecular patterns are affected by mutations in *STT3A* (Fig. 1, Supplementary Fig. S1).

In this study, we demonstrated that *STT3A* is involved in the recognition of sphingolipids and affects the molecular mass of NCER2. In addition, *stt3a* mutants showed a reduced response to 9-methyl-branched sphingoid base, suggesting that *STT3A* might also regulate RDA2 or its downstream signaling factors. Comparative glycoproteomics between WT and *stt3a* plants should help reveal the importance of N-glycosylation in PTI signaling.

## MATERIALS AND METHODS

### Plant materials and growth conditions

All Arabidopsis (*Arabidopsis thaliana*) lines used in this study were in the Col-0 background. Seeds of the *stt3a-2* (SALK_058814) and *rda2-3* (SALK_143489C) mutants were obtained from the Arabidopsis Biological Resource Center (ABRC). The *rda2-5* and *ncer2-2* mutants were described previously (Kato et al., 2022). All seeds were surface sterilized using a sodium hypochlorite solution and incubated at 4°C in the dark for 4 days prior to seed sowing. Seeds were plated on Murashige and Skoog (MS) (Murashige and Skoog, 1962) liquid medium containing 0.5% (w/v) sucrose, B5 vitamin solution, and 2 mM MES (pH 5.7) and incubated at 23°C in the light (24-h photoperiod, 57 μmol m^-2^ s^-1^). For seed propagation, Arabidopsis plants were grown on MS agar medium (MS salt, 1.5% sucrose, B5 vitamin solution, 2 mM MES [pH 5.7], and 0.8% agar) for 10 to 14 days, transferred to soil, and growth under continuous light at 23°C (57 μmol m^-2^ s^-1^). Soil-grown Arabidopsis plants were grown under controlled conditions (23°C, 10-h photoperiod, 57 μmol m^-2^ s^-1^)

### Genotyping T-DNA insertion lines

Genomic DNA was extracted from seedlings in sucrose lysis buffer containing 50 mM Tris-HCl (pH 8.0), 300 mM NaCl, and 300 mM sucrose. PCR products were obtained with KOD One PCR Master Mix (TOYOBO) and separated by electrophoresis on 1% (w/v) agarose gels. Left primer (LP) and border primer (BP) were used as forward primers, and right primer (RP) was used as the reverse primer (Supplementary Table S2) to identify lines homozygous for the respective T-DNA insertion. The PCR conditions were 98°C for denaturation, 55°C for annealing, and 68°C for extension.

### Elicitors and chemicals

Pi-Cer D was purified from *Phytophthora infestans* as described by Monjil et al. (2024). Other elicitors and chemicals were acquired or purchased from commercial sources: flg22, elf18, and nlp24 (Life Technologies, Tokyo, Japan); chitin (C9752, Sigma); (4E,8E)-9-methyl-4,8-sphingadienine (9Me-Spd, NS440901; Nagara Science); 3-hydroxy decanoic acid (3-OH-FA, 24613; Cayman). Cello-oligosaccharides (COS) were prepared via hydrolysis of cotton linters or microcrystalline cellulose (Avicel PH 101, Sigma-Aldrich, Burlington, MA, USA).

### Measurement of bioluminescence responses

Surface-sterilized seeds were incubated at 4°C in the dark for 4 days and sown in the wells of 96-well microplates (Luminunc^TM^ Plates White F96; Thermo Fisher Scientific) containing 150 µl liquid MS medium supplemented with 50 µM D-luciferin, potassium salt (Biosynth). Seed germination was conducted under continuous illumination (23°C, 57 μmol m^-2^ s^-1^). Following a 7-day incubation period, the seedlings were treated with elicitors (100-fold dilution), and the 96-well plates were sealed with a plate seal (100-THER-PLT, Excel Scientific) instead of a plastic cover. Bioluminescence from each well was measured automatically and immediately after elicitor addition using a bioluminescence monitoring system (model CL96-4; Churitsu Electric Corp.) with a robotic plate conveyor (model CI-08L; Churitsu Electric Corp.). Bioluminescence data were analyzed using the software provided with the instrument (SL00-01; Churitsu Electric Corp.).

### Generation of F_2_ progeny and whole-genome sequencing

F_1_ progeny were generated by crossing the p*WRKY33-LUC* reporter line (W33-1B, the parental line of the mutants) (Kato et al., 2022) with each of the mutants identified in the screen. All F_1_ plants were subsequently self-pollinated to obtain F_2_ seeds. The bioluminescence responses of the F_2_ seedlings were tested using Pi-Cer D treatment as described above. F_2_ seedlings with the mutant phenotype were selected and their genomic DNA extracted. The same quantity of genomic DNA from 30 F_2_ seedlings with the mutant phenotype was combined to obtain a bulk (pooled) DNA sample for MutMap analysis. For whole-genome sequencing, genomic DNA samples were extracted from young (3-week-old) leaves using a DNeasy Plant Mini Kit (Qiagen). Sequencing libraries were prepared for the L-08, L-13, and L-51 mutant lines using an Illumina TruSeq DNA LT Sample Prep Kit (Illumina). All libraries were sequenced as paired-end 150-bp reads. The libraries were sequenced on a HiSeq High-Output system. The whole-genome sequencing data have been deposited in DDBJ BioProject under accession number PRJDB20249.

### MutMap analysis

The p*WRKY33-LUC* reporter line reference sequence was constructed by replacing nucleotides in Col-0 with those of the p*WRKY33*-*LUC* reporter line W33-1B (Kato et al. 2022). Bulked segregant analysis, as implemented in MutMap (Abe et al., 2012), was performed using the MutMap pipeline (https://github.com/YuSugihara/MutMap) (Sugihara et al., 2022). The Arabidopsis Col-0 reference genome was downloaded from ftp://ftp.ensemblgenomes.org/pub/release-36/plants/fasta/Arabidopsis_thaliana/dna/Arabidopsis_thaliana.TAIR10.dna.toplevel.fa.

Whole-genome sequencing of the three mutants identified 773 ± 334 (mean ± s.d.; range 398– 1,038) SNPs relative to the p*WRKY33-LUC* reporter parental line W33-1B.

### Plasmid construction and plant transformation

To complement the *stt3a* mutants (*stt3a-3*, *stt3a-4*, *stt3a-5*), a 10.7-kb genomic fragment containing the *STT3A* genomic coding region and its native promoter was amplified and inserted into pBIB-KAN at the SalI and SacI restriction sites using an In-Fusion HD Cloning Kit (Takara). The primer sequences used for plasmid construction are listed in Supplementary Table S2.

The plasmids were introduced into Agrobacterium (*Agrobacterium tumefaciens*) strain GV3101::pMP90 by electroporation and used to transform Arabidopsis plants via the floral dip method (Clough and Bent, 1998). T_1_ seeds were collected from these T_0_ plants, and T_1_ seedlings were grown on MS medium containing kanamycin (50 μg/ml) and carbenicillin (150 μg/ml) to select transgene-positive T_1_ plants. T_2_ seeds were then collected from the positive T_1_ plants and sown on MS medium containing kanamycin; T_2_ lines showing a 3:1 segregation ratio were thought to contain a single T-DNA. T_3_ plants homozygous for the transgene were obtained and used for subsequent analysis.

### MAPK activation assay

Seedlings germinated from surface-sterilized seeds were grown in 96-well microplates in MS liquid medium for 7 days. Samples were obtained from sphingolipid-treated seedlings at 15 and 30 min of treatment (0.17 µM Pi-Cer D, 0.5 µM 9Me-Spd) using the same method used for screening. All samples were immediately frozen in liquid nitrogen. Proteins were extracted from the samples in extraction buffer (50 mM HEPES, pH 7.4, 5 mM EDTA, 0.5 mM EGTA, 50 mM β-glycerophosphate, 10 mM Na_3_VO_4_, 10 mM NaF, 2 mM DTT). MAPK activation was monitored by immunoblot analysis using an antibody that recognizes the dual phosphorylation of the activation loop of MAPK (pTEpY). Phospho-p44/42 MAPK (Erk1/2; Thr-202/Tyr-204, Cell Signaling Technology), and rabbit monoclonal antibodies were used to detect phosphorylated MAPK according to the manufacturer’s protocol (#9101, Cell Signaling Technology). Immunoblotting detection solution (ECL Prime Western Blotting Detection Reagents, Amersham) was used for signal detection. The PVDF membrane was stained with Coomassie Brilliant Blue (PageBlue Protein Staining Solution, Thermo Fisher Scientific) to verify equal protein loading.

### RNA extraction and RT-qPCR

Seven-day-old seedlings were treated with sphingolipids, and samples were collected at the specified time points before being frozen in liquid nitrogen. Total RNA was extracted from the samples using an RNeasy Plant Mini Kit (Qiagen) according to the manufacturer’s instructions. Following RNA extraction, first-strand cDNA was synthesized using Takara PrimeScript RT Master Mix (Takara). Quantitative PCR was conducted using rTaq DNA polymerase (TAP201, TOYOBO) and EvaGreen Dye (31000, Biotium, Inc.) on a CFX Connect Real-Time System (BIO-RAD) with the primers listed in Supplementary Table S2. Relative gene expression levels were calculated using the ΔΔCt method, with *ubiquitin-conjugating enzyme 21* (*UBC21*, At5g25760) (Czechowski et al., 2005) serving as a reference for normalization.

### Pathogen inoculation assay

The inoculation assay was performed as previously reported (Singkaravanit-Ogawa et al., 2021) with some modifications. *Colletotrichum higginsianum* strain MAFF_305635 was cultured on potato dextrose agar (BD Biosciences, Franklin Lakes, NJ) at 24°C in the dark. For plant inoculation, four drops (5 µl per drop) of a spore suspension containing 1.5 × 10^5^ conidiospores/ml were placed onto one leaf of a 5-to 6-week-old Arabidopsis plant without punching. The plants were covered with a transparent lid to maintain high humidity conditions and placed in a growth cabinet at 23°C under a 10-h light/14-h dark photoperiod until the day of observation (4 dpi). The inoculated leaves were excised and photographed after 4 days for analysis. The lesion size (area) was quantified using ImageJ (https://imagej.net/ij/). The statistical significance of differences between means was determined using Tukey’s test. In the figures, different lowercase letters indicate significant differences (*p* < 0.05).

### Production of the anti-NCER2 antibody

A peptide from the C-terminus of NCER2 (ADVPPKSTFRR) with an N-terminal cysteine was chemically synthesized (Scrum). The antigen peptide was conjugated to a keyhole limpet hemocyanin carrier via SS-linkage through the N-terminal cysteine, and polyclonal antisera were raised in rabbits (Scrum). Anti-NCER2 antibodies were purified from the antisera by affinity chromatography using antigen-immobilized beads (HiTrap NHS-activated HP; GE Healthcare).

### Extraction of apoplast wash fluid and detection of NCER2

Apoplast wash fluid (AWF) was extracted from the samples as described by Gentzel et al. (2019), with minor modifications. Extraction buffer (20 mM Tris-HCl, pH 7.5, 50 mM NaCl) was introduced into leaves via syringe infiltration. The leaves were blotted dry with a paper towel, wrapped around a 1-ml pipette tip using Parafilm, and placed in a 15-ml conical tube. The tube was subjected to centrifugation at 1,000 × *g* for 3 min at room temperature using a swing rotor. AWF proteins were precipitated by acetone precipitation and resuspended in extraction buffer containing 1% SDS. The protein content of the AWF was quantified using the Bradford method, and a 10 µg equivalent of AWF proteins was separated by electrophoresis on a 12% (w/v) SDS acrylamide gel. Immunoblot analysis was performed with anti-NCER2 antibodies (1:2,000 dilution) and HRP-conjugated secondary antibodies (1:2,000 dilution).

### Accession numbers

The accession numbers of the Arabidopsis genes mentioned in this article are as follows: *STT3A* (At5g19690), *RDA2* (At1g11330), *NCER2* (At2g38010), and *WRKY33* (At2g38470).

## ACKNOWLEDGEMENTS

We thank Y. Nakagawa (Kyoto University) for technical support. We also thank Prof. Kazuhito Kawakita and Makoto Ojika (Nagoya University, Japan) for valuable suggestions and for Pi-Cer D purification. Some of the computational analysis was performed on the National Institute of Genetics (NIG) supercomputer at the Research Organization of Information and Systems (ROIS), NIG, Tokyo, Japan.

## AUTHOR CONTRIBUTIONS

Conceptualization: SC, RT, HK

Methodology: MS, AA, RT, HK

Investigation: SC, NI, YI

Visualization: SC, MS, AA

Funding acquisition: YT, RT, HK

Project administration: DT, YT, RT, HK

Supervision: DT, KS, YT, RT, HK

Writing – original draft: SC

Writing – review & editing: SC, DT, YT, KS, RT, HK

## Supplementary Information

**Supplementary Fig. S1.**
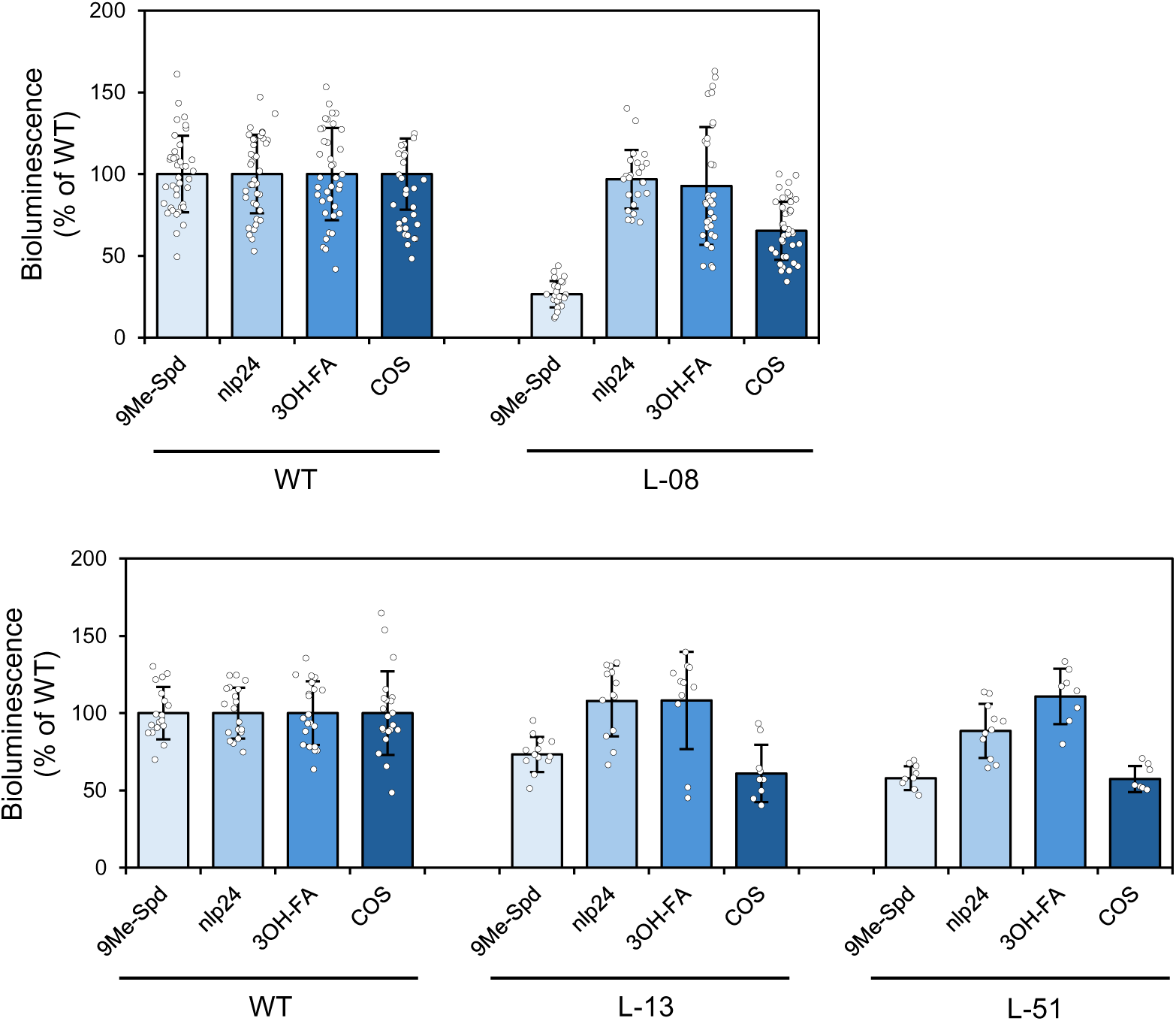
Bioluminescence responses of the three mutants to treatment with four elicitors. Seven-day-old wild type (p*WRKY33-LUC* reporter line; WT), L-08, L-13, and L-51 seedlings were treated with 0.5 µM (4E,8E)-9-methyl-4,8-sphingadienine (9Me-Spd), 0.1 µM nlp24, 1 µM 3-hydroxy fatty acid (3-OH-FA), or 20 µg/ml cello-oligosaccharides (COS). The bioluminescence level of the WT was set to 100%, to which the bioluminescence levels in the mutants were normalized.

**Supplementary Fig. S2.**
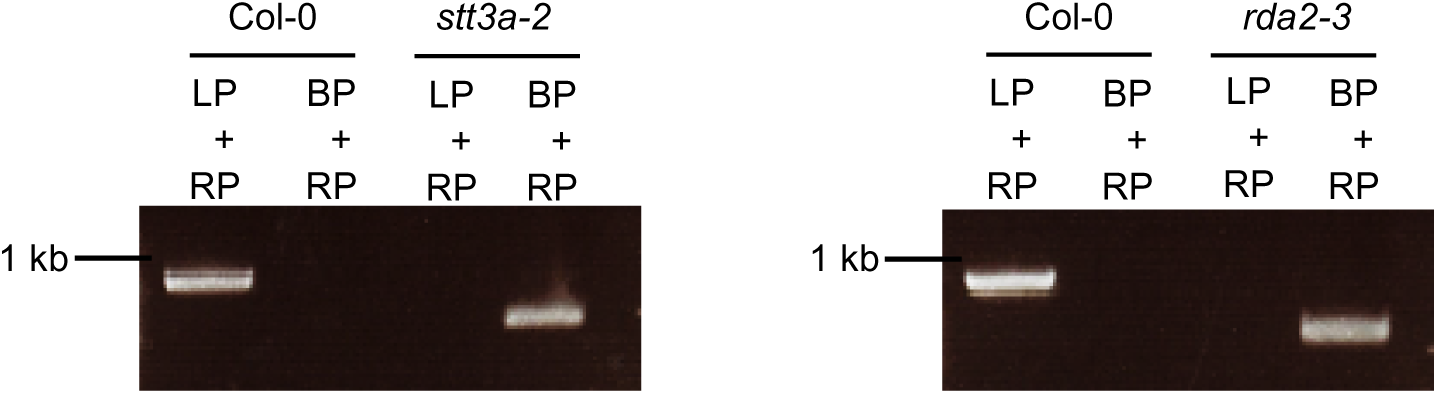
Confirmation of the T-DNA insertion in the *stt3a-2* and *rda2-3* mutant lines. Gel electrophoresis of the PCR products confirmed the genotypes of the T-DNA insertion lines and Col-0. LP and RP indicate the left primer and right primer of the respective gene, respectively; BP indicates the T-DNA border primer.

**Supplementary Fig. S3.**
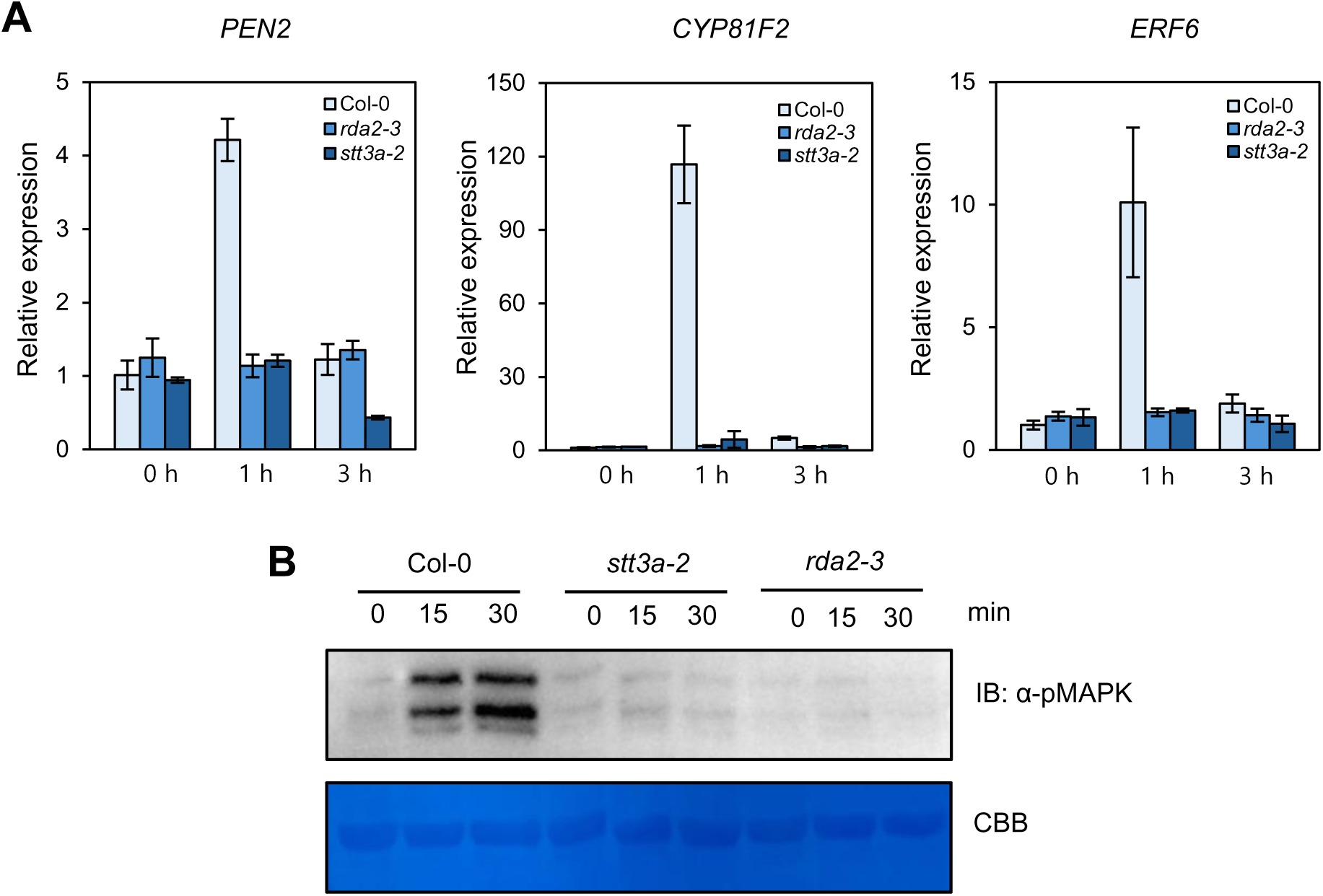
Immune response analysis in the *stt3a* T-DNA insertion mutant treated with Pi-Cer D. (A) Relative expression levels of the defense-related genes *PEN2*, *CYP81F2*, and *ERF6* as determined by RT-qPCR. Seven-day-old Col-0, *stt3a-2*, and *rda2-3* seedlings were treated with 0.17 µM Pi-Cer D, and sampled at 0, 1, or 3 h after treatment (*n* = 3). (B) MAPK phosphorylation assay in Col-0, *stt3a-2*, and *rda2-3.* Seven-day-old seedlings were sampled at 0, 15, or 30 min of treatment with 0.17 µM Pi-Cer D. Phosphorylated MAPKs were visualized with anti-phospho-p44/p42 MAPK antibody. Equal protein loading was determined by staining the membrane with Coomassie brilliant blue (CBB).

**Supplementary Fig. S4.**
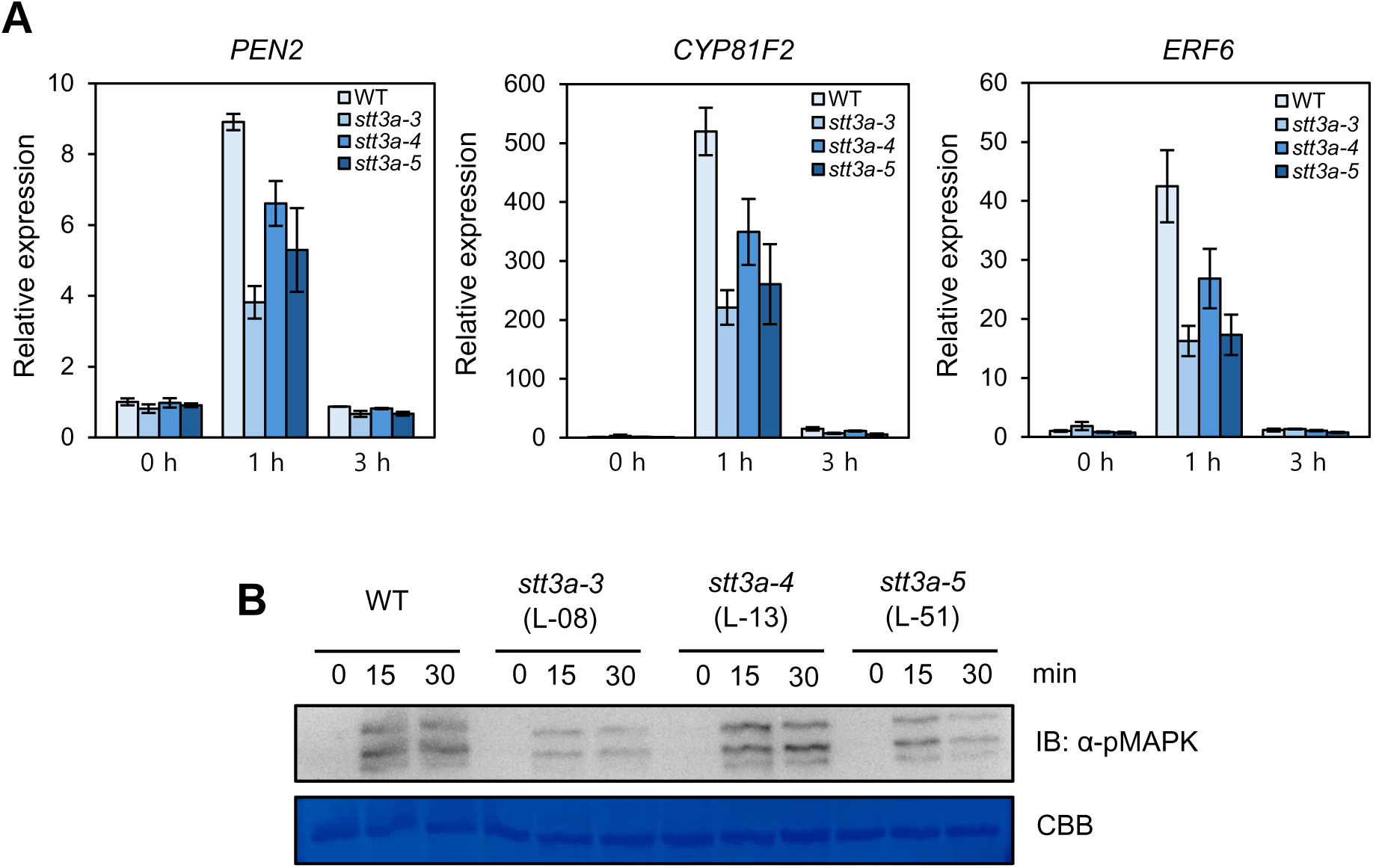
Immune response analysis in the *stt3a-3, stt3a-4, and stt3a-5* mutants treated with 9Me-Spd. (A) Relative expression levels of the defense-related genes *PEN2*, *CYP81F2*, and *ERF6* as determined by RT-qPCR (means ± SE). Seven-day-old seedlings of the p*WRKY33-LUC* reporter line (WT), *stt3a-3* (L-08), *stt3a-4* (L-13), and *stt3a-5* (L-51) were treated with 0.5 µM 9Me-Spd, and sampled at 0, 1, and 3 h after treatment (*n* = 3). (B) MAPK phosphorylation assay in WT and each *stt3a* mutant. Seven-day-old seedlings were sampled at 0, 15, and 30 min of treatment with 0.5 µM 9Me-Spd. Phosphorylated MAPKs were visualized with anti-phospho-p44/p42 MAPK antibody. Equal protein loading was determined by staining the membrane with CBB.

**Supplementary Fig. S5.**
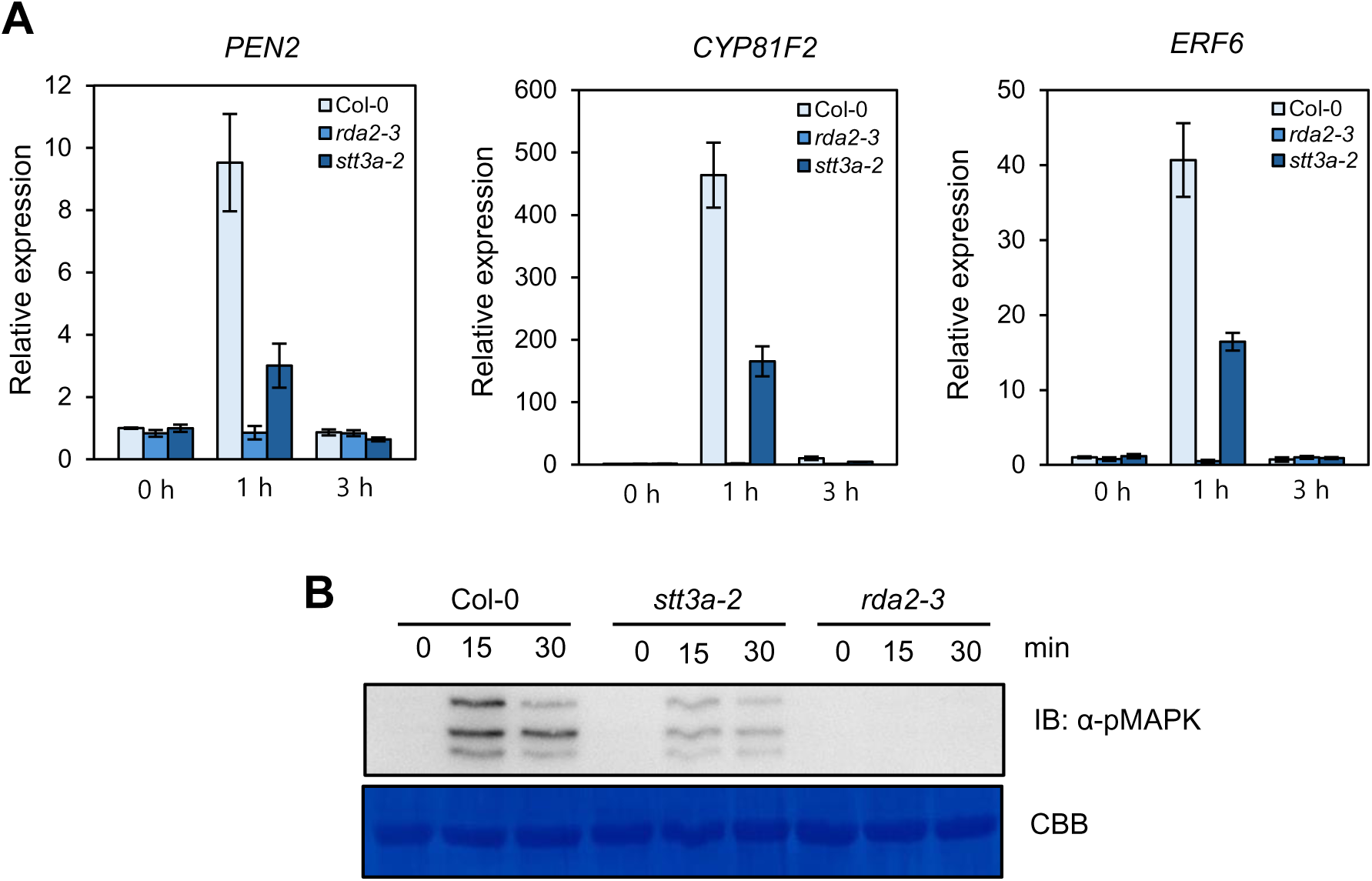
Immune response analysis in the *stt3a* T-DNA insertion mutant treated with 9Me-Spd. (A) Relative expression levels of the defense-related genes *PEN2*, *CYP81F2*, and *ERF6* as determined by RT-qPCR. Seven-day-old seedlings of Col-0, *stt3a-2*, and *rda2-3* were treated with 0.5 µM 9Me-Spd, and sampled at 0, 1, and 3 h after treatment (*n* = 3). (B) MAPK phosphorylation assay in Col-0, *stt3a-2*, and *rda2-3.* Seven-day-old seedlings were sampled at 0, 15, and 30 min of treatment with 0.5 µM 9Me-Spd. Equal protein loading was determined by staining the membrane with CBB.

---

**Supplementary Table S1.**
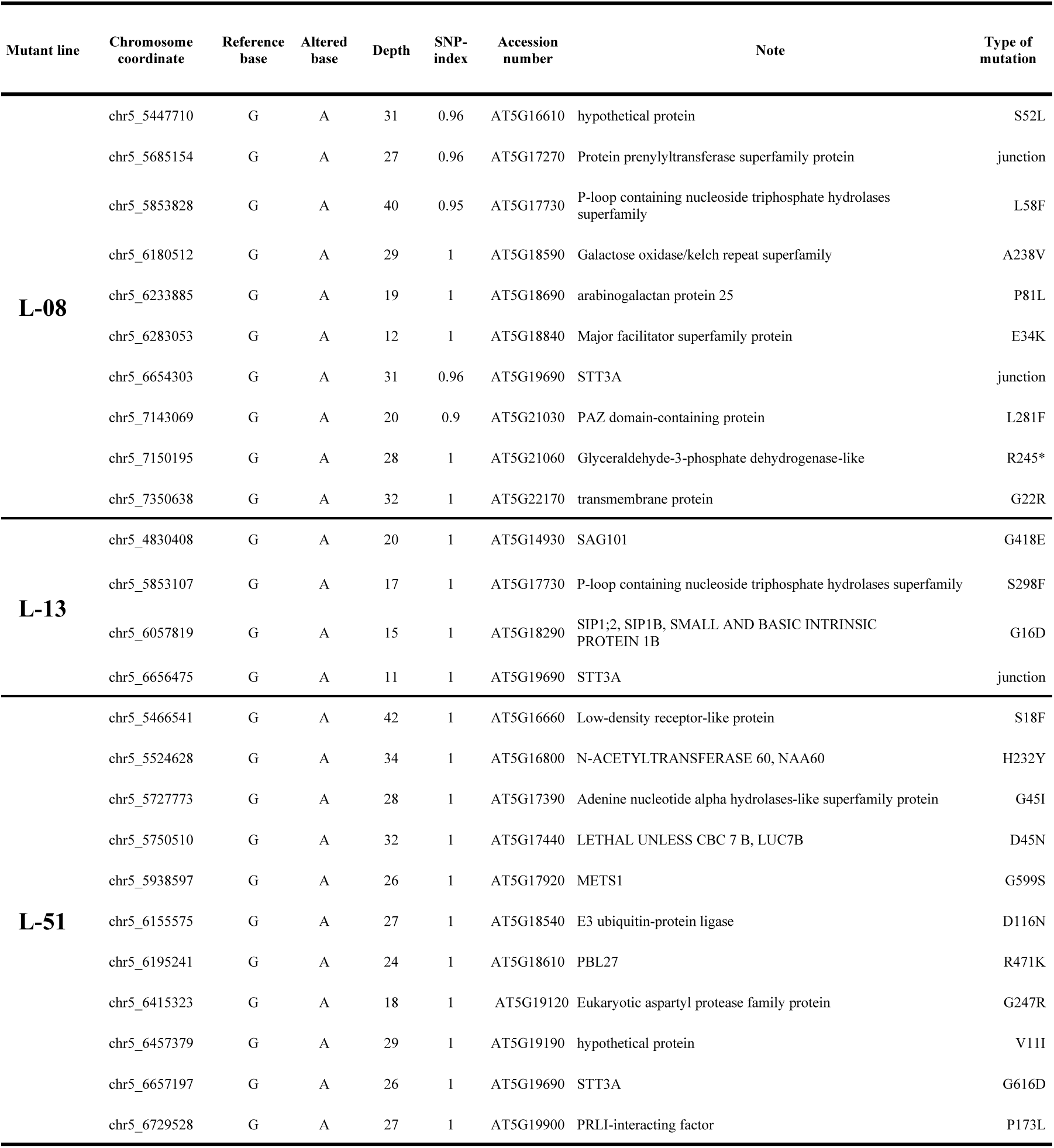
Candidate genes and polymorphisms identified in the high SNP-index intervals on Arabidopsis chromosome 5.

**Supplementary Table S2.**
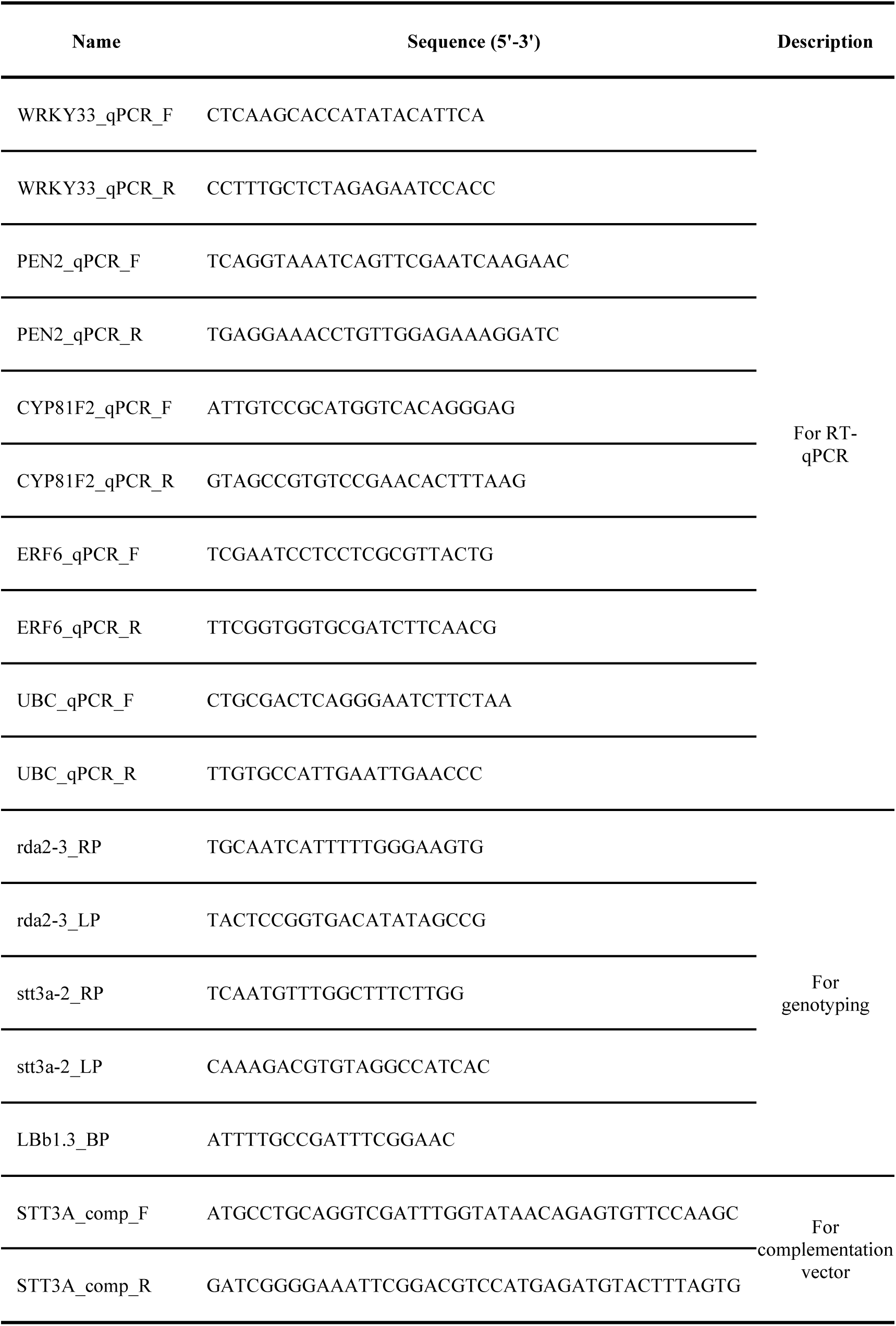
Primers used in this study.

